# *Aedes aegypti* SNAP and a calcium transporter ATPase influence dengue virus dissemination

**DOI:** 10.1101/2020.07.10.198226

**Authors:** Alejandro Marin-Lopez, Junjun Jiang, Yuchen Wang, Yongguo Cao, Tyler MacNeil, Andrew K Hastings, Erol Fikrig

## Abstract

Dengue virus (DENV) is a flavivirus that causes marked human morbidity and mortality worldwide, being transmitted to humans by *Aedes aegypti* mosquitoes. Habitat expansion of *Aedes*, mainly due to climate change and increasing overlap between urban and wild habitats, places nearly half of the world’s population at risk for DENV infection. After a bloodmeal from a DENV-infected host, the virus enters the mosquito midgut. Next, the virus migrates to, and replicates in, other tissues, like salivary glands. Successful viral transmission occurs when the infected mosquito takes another blood meal on a susceptible host and DENV is released from the salivary gland via saliva into the skin. During viral dissemination in the mosquito and transmission to a new mammalian host, DENV interacts with a variety of vector proteins, which are uniquely important during each phase of the viral cycle. Our study focuses on the interaction between DENV particles and protein components in the *A. aegypti* vector. We performed a mass spectrometry assay where we identified a set of *A aegypti* salivary gland proteins which potentially interact with the DENV virion. Using dsRNA to silence gene expression, we analyzed the role of these proteins in viral infectivity. Two of these candidates, a synaptosomal-associated protein (AeSNAP) and a calcium transporter ATPase (ATPase) appear to play a role in viral replication both *in vitro* and *in vivo*. These findings suggest that AeSNAP plays a protective role during DENV infection of mosquitoes and that ATPase protein is required for DENV during amplification within the vector.

**Importance:** *Aedes aegypti* mosquitoes are the major vector of different flaviviruses that cause human diseases, including dengue virus. There is a great need for better therapeutics and preventive vaccines against flaviviruses. Flaviviruses create complex virus-host and virus-vector interactions. The interactions between viral particles and protein components in the vector is not completely understood. In this work we characterize how two mosquito proteins, “AeSNAP” and “ATPase”, influence DENV viral dissemination within *A. aegypti*, using both *in vitro* and *in vivo* models. These results elucidate anti-vector measures that may be potentially be used to control dengue virus spread in the mosquito vector.

## Introduction

Dengue is a major public health threat in tropical and subtropical areas, and as climate change and urbanization continues, the illness may spread to other locations across the globe (1). According to WHO reports, before 1970 only nine countries experienced outbreaks of severe dengue. Today, the disease is endemic in more than 100 countries in Africa, the Americas, South-East Asia, the Western Pacific regions, and the Eastern Mediterranean regions. Recently, some dengue cases have been documented in places where the disease was absent for more than 50 years, including France and Spain (European Centre for Disease Prevention and Control). Therefore, more attention is required to counteract this expansion, fed by processes including global warming, unprecedented human mobility, rapid urban population growth, and large-scale changes in ecosystems (2–5).

Dengue virus (DENV) is a positive-sense, single-stranded RNA virus that belongs to the genus *Flavivirus* within the family *Flaviviridae*. Its primary vector is the *Aedes aegypti* mosquito. After taking a viremic blood meal, DENV establishes infection in the midgut. The midgut represents the first barrier to block viral propagation in the mosquito. Upon establishing a successful infection, the virus disseminates systemically through the hemolymph where it can invade secondary tissues, such as the salivary glands (6). Replication in the salivary glands leads to virion release into the saliva, the last step prior to virus transmission to the a human host (7).

Dengue is provoked by four serologically different DENV serotypes and usually results in a mild self-limiting disease, but is also capable of causing much more severe dengue hemorrhagic fever (DHF) or dengue shock syndrome (DSS) with approximately 20,000 fatalities recorded annually (8). Conventional vaccines are in development and some of them are being implemented in a few countries (9). The development of these vaccines has been complicated by to the co-circulation of different serotypes and the phenomena of antibody-dependent enhancement (ADE) (10–13), ADE occurs when an individual who has encountered one DENV serotype is infected by a second DENV serotype and non-neutralizing antibodies bind to the virus allowing it to enter mononuclear cells, susceptible to virus infection, via an FcR-dependent mechanism. This process leads to greatly enhanced disease severity (14). Therefore, exploring novel methods to block DENV spread in the mosquito vector by analyzing ways to interfere with vector proteins-virus interaction, could be a good alternative to, or complement for conventional vaccines. The targeting, modification, or elimination of specific genes in *A. aegypti* can reduce vector competence for virus acquisition, dissemination, and transmission (15–17), reducing the expansion of this widespread arboviral disease. Indeed, transmission-blocking vaccines (TBV) may trigger a strong immune response from a vertebrate host against mosquito components which can regulate the viral infection in vector tissues and lead to the blockage of viral infection in the vector (18). This has been shown to be the case for C-type lectins and the cysteine rich venom protein CRVP379 in the mosquito. When the interaction between the virus and vector proteins are blocked using specific antibodies, DENV infection in *A. aegypti* is effectively interrupted (19, 20).

Dengue replication in the salivary gland is the last step before virus transmission to the host, and little is known about the protein interactions that take place at this stage. Here, we explore the impact of altering protein expression levels of several *A. aegypti* proteins found in the salivary gland tissue during DENV infection, *in vitro* and *in vivo*. Using viral purification coupled to a mass spectrometry assay, we identified a set of *A. aegypti* salivary gland proteins which potentially interact with DENV virions. Next, we used dsRNA silencing to analyze the effect of these interaction candidates during DENV infection. Using these techniques, we demonstrate that a synaptosomal-associated protein, that we named here AeSNAP, and a calcium transporter ATPase protein (ATPase) have a role in DENV infection *in vitro*, in the Aag2 *A. aegypti* cell line, and in *vivo* in the *A. aegypti* mosquito. Silencing of AeSNAP expression led to an increase in viral burden at 24 hours post-infection (hpi) *in vitro* and 7 dpi in the mosquito, whereas we found the opposite result after silencing ATPase protein expression. These findings suggest that AeSNAP may have a protective role during DENV infection whereas ATPase protein is required for DENV during amplification. This highlights two possible targets for controlling DENV transmission in the mosquito vector.

## Results

### Identification of DENV binding proteins in *A. aegypti* salivary glands using a mass spectrometry assay

To identify *A. aegypti* salivary gland components that potentially interact with DENV virions, we utilized gradient sucrose purification of DENV virions that were pre-incubated with salivary gland extracts (SGE). A DENV only was used as control, we were able to identify vector peptides that were only detected in samples containing SGE. Using the National Center for Biotechnology Information bioinformatic search database (BLASTp), we identified peptides that were conserved in *A. aegypti*, but not *A. albopictus*, resulting in 45 *A. aegypti* salivary gland proteins that potentially interact with DENV virions. A list of these putative DENV binders obtained in three different runs was assembled (Table 1) and a Venn diagram was generated displaying the number of hits in each biological replicate and the overlap between the three runs (Figure 1). Eight, 19, and 38 proteins were found in runs 1, 2 and 3 respectively. We identified two unique proteins in only the 1^st^ and the 3^rd^ runs, eight unique proteins in only the 2^nd^ and the 3^rd^ runs and no overlapping proteins in only the 1^st^ and 2^nd^ runs. Finally, we identified five unique proteins in all three runs. The subset of the proteins which overlapped in multiple runs were then analyzed in subsequent experiments for their effect on DENV infection.

**Figure 1.**
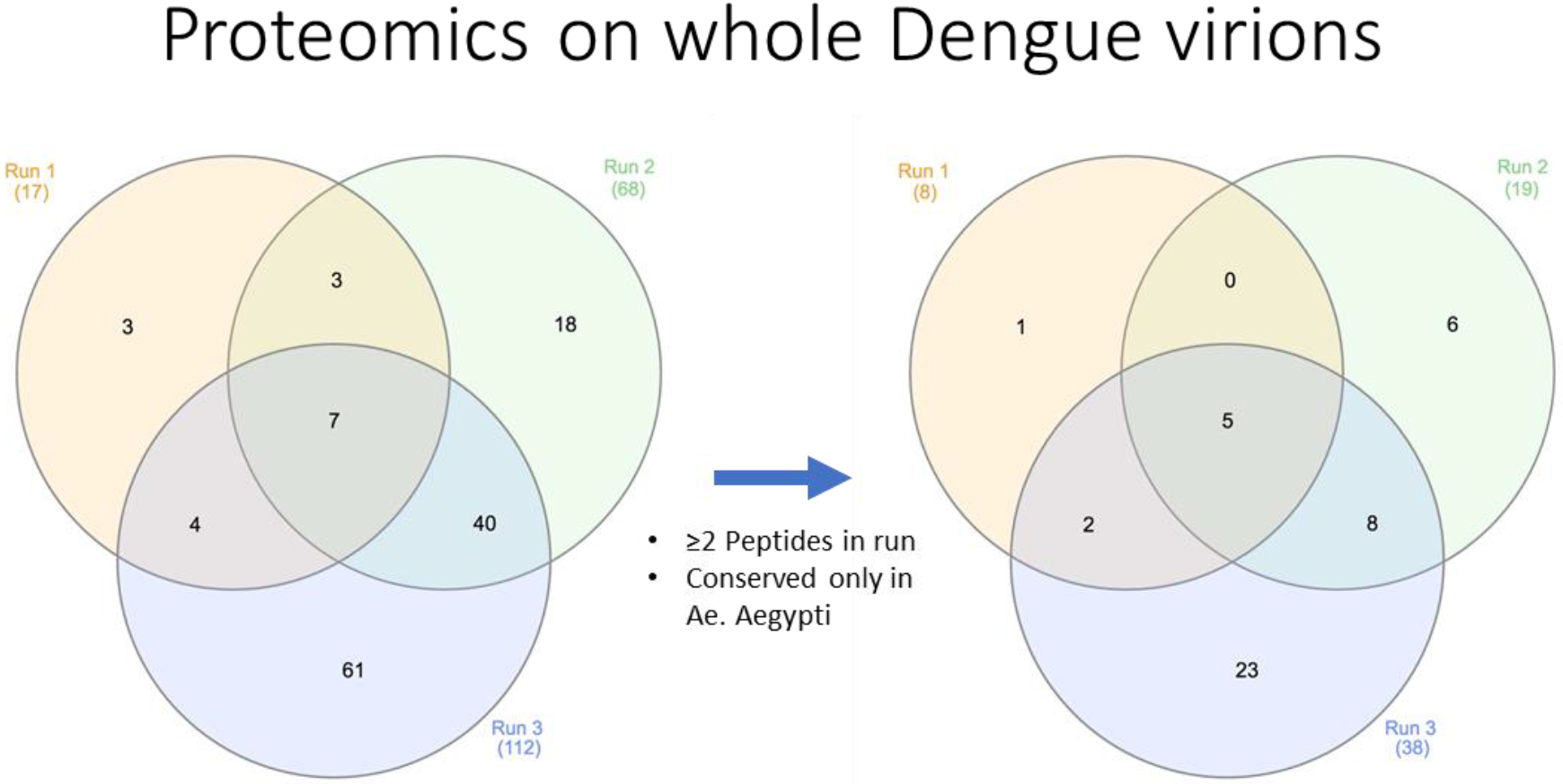
Illustration of a three-cycle Venn diagram with the hits recovered from the mass spectrometry assay before selection (left) and after selection (right) of the hits. Selection was based on the number of sequences found for every hit and their conservation in *A aegypti*.

**Table 1.**
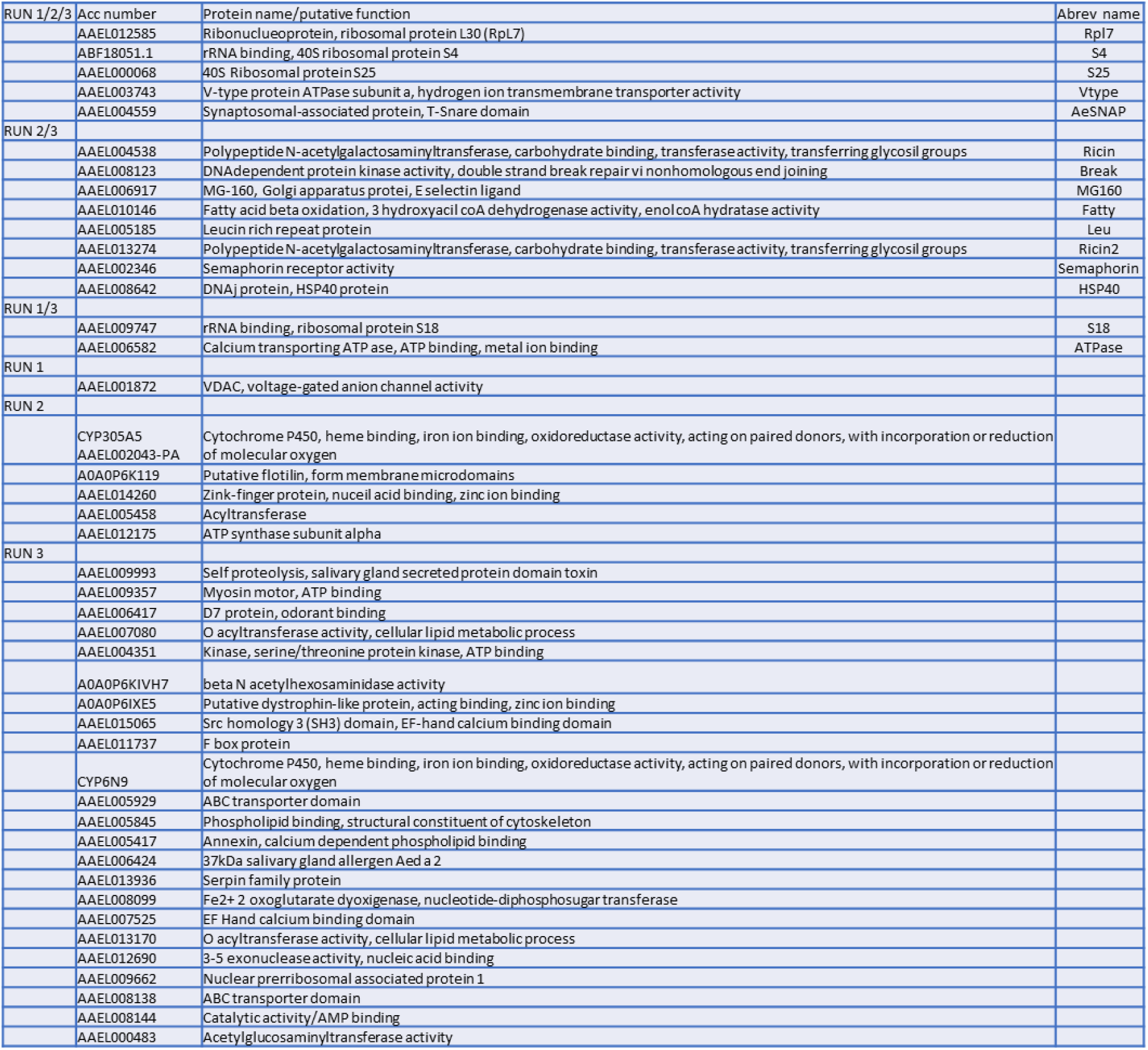
List of putative DENV binders obtained from the *A aegypti salivary* gland extract by mass spectrometry assay. Three different experiments of mass spectometry assays were performed, and hits were clustered according to their presence in every experiment. Accession numbers, putative functions and abbreviated names were shown.

### Silencing genes which encode salivary gland proteins associated with virions, alters DENV infection in a mosquito cell line

To elucidate the role of the protein candidates obtained from our mass spectrometry analysis (shown in Table 1) during DENV infection, we used RNAi to reduce gene expression and analyzed the effect on viral infection. dsRNAs were generated against the genes encoding proteins that were found in at least 2 runs of the mass spectrometry analysis and used to silence these genes in an *Ae*. *aegypti* cell line, Aag2. A reduction between 75% and 95% in the mRNA transcripts was achieved at 48 hours post transfection in the genes screened, analyzed by qRT-PCR, with two exceptions (MG160 and Break), which were removed for further analysis (Fig 2).

**Figure 2.**
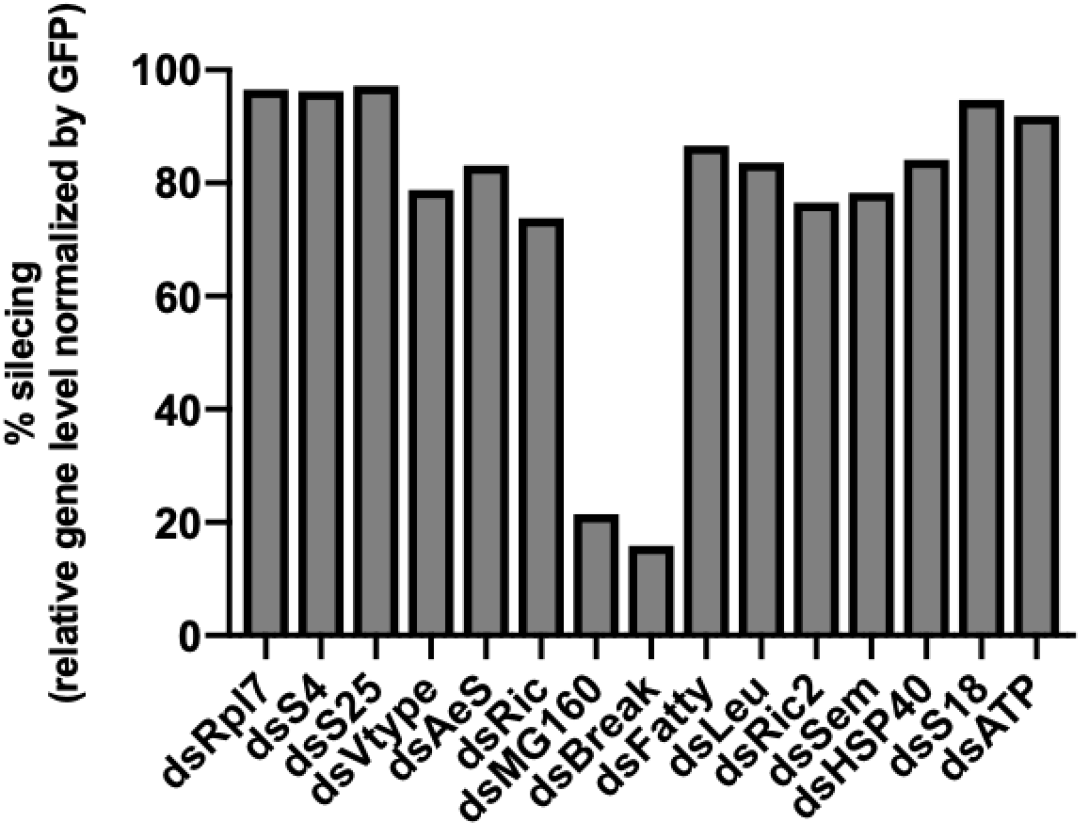
dsRNA silencing efficacy of A. aegypti genes in Aag2 cells. Hits from at least two experiments listed in table 1 were knocked down in Aag2 cells using RNAi. At 48 h post-knockdown, silencing efficiency was analyzed by qRT-PCR, obtaining the relative levels of the specific gene normalized by Rp49 as housekeeping. Data is displayed as knockdown percentage of every hit compared to control (dsGFP). qRT-PCR analysis was done in pentaplicate, and the percentage of silencing was obtained comparing mean values of the relative gene levels between the specific genes and the GFP control (100%).

To identify the effect of protein knockdown on DENV infection, Aag2 cells were transfected with specific dsRNAs and 48 hours later were infected with DENV2 (MOI of 1.0). Each sample was then analyzed for intracellular viral production using qRT-PCR at 6, 9, 12 and 24 h post-infection (Fig 3). Knockdown of genes encoding several proteins led to significant changes in the intracellular viral load. At 6, 9 and 12 h post-infection, DENV titer was reduced in AeSNAP and Ric silenced cells as well as S18 at 9 and 12 hpi and Ric2, Sem, HSP40 and ATPase expression levels at 24 hpi. In contrast, we observed a significant increase in the DENV viral burden at 24 hpi for Vtype and surprisingly AeSNAP silenced cells (Fig 3).

**Figure 3.**
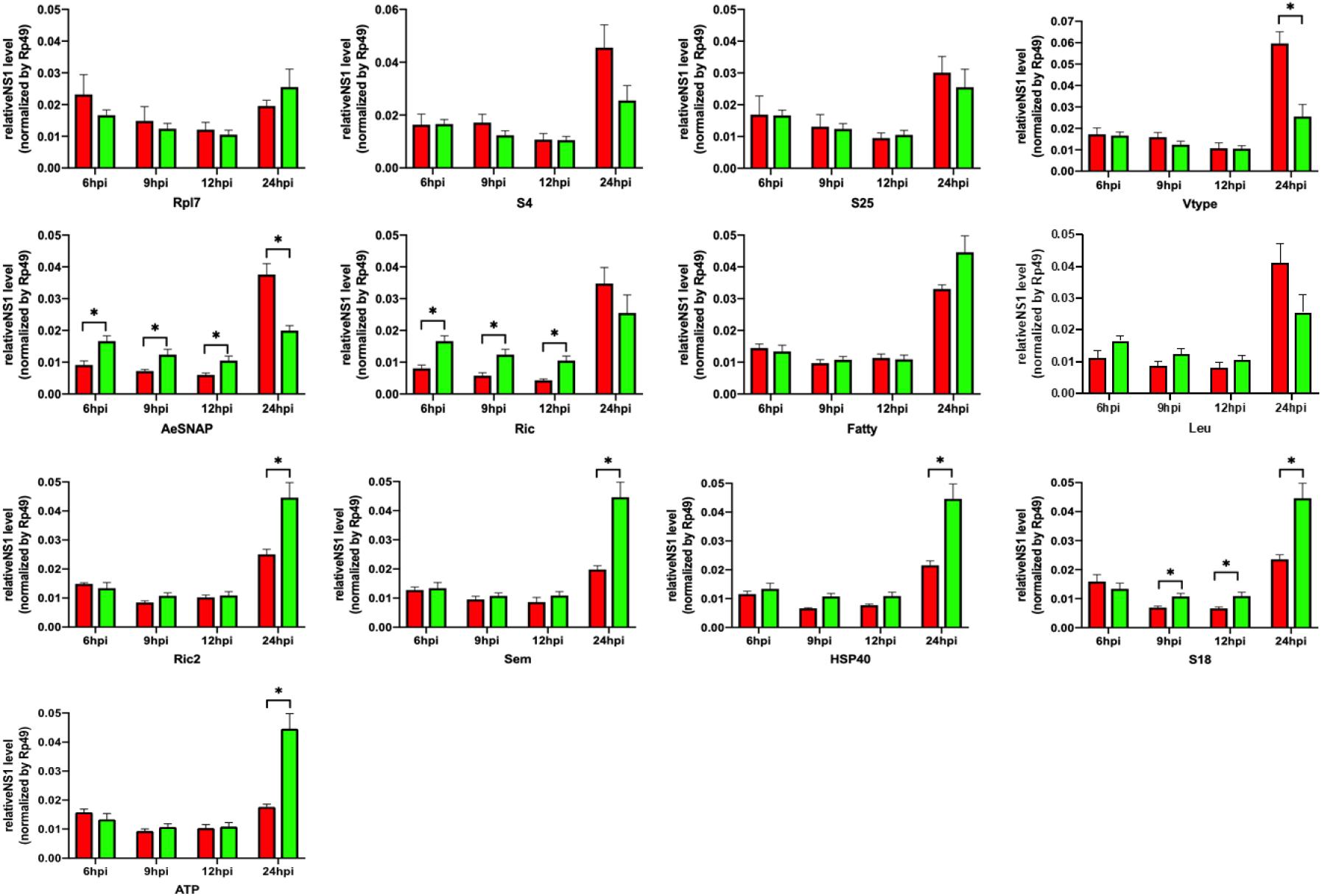
DENV infection relative levels in Aag2 cells. Viral burden was analyzed in Aag2 cells infected with DENV2 (MOI of 1.0) and was measured using qRT-PCR analysis at the timepoints indicated. Samples were taken at 6, 9, 12 and 24h post-knockdown to see the effect of silencing during DENV2 infection. The results represent the averages from samples done in pentaplicate, with the mean and standard deviation. In green, GFP-silenced control cells. In red, protein-silenced cells. Asterisks represent significant difference between samples, calculated by the Mann-Whitney nonparametric test (*P* < 0.05).

### Silencing AeSNAP and ATP proteins alters DENV dissemination in *A. aegypti* mosquitoes

After the *in vitro* analysis, we focused on two of the vector proteins that showed the greatest ability to alter viral replication, AeSNAP and ATPase proteins. The AeSNAP protein is a member of the Soluble N-ethylmaleimide-Sensitive Factor Attachment Protein (SNAP) protein family. This protein was particularly interesting, because SNAPs are known mainly to be involved in vesicle trafficking. The ATPase protein belongs to the calcium transporter ATPase proteins, major players in maintaining calcium homeostasis in the cell. Therefore, we decided to assess the role of AeSNAP and ATPase proteins during DENV dissemination in the mosquito vector. For this aim, we intrathoracically injected *A. aegypti* mosquitoes with one μg of AeSNAP or ATPase dsRNA, and, 72 hours post injection, injected these same mosquitoes intrathoracically again with 100 PFU of DENV2. DENV titer was then analyzed at 4 or 7 dpi (7 and 10 days post dsRNA injection, respectively) (Fig 4A). To confirm AeSNAP gene knockdown, silencing efficiency was checked at these timepoints, and a significant reduction in the AeSNAP RNA transcript level (red) compared to the GFP control group (green) was observed (Fig 4B). Finally, DENV viral burden was analyzed, observing a tendency in viral burden increase at 4 dpi and a significant increase in the AeSNAP knockdown group (red) at day 7 post-infection. (Fig 4C). In addition, we also analyzed the role of ATPase during DENV dissemination. *Aedes* mosquitoes were silenced with ATP dsRNA (purple), and the silencing efficacy was confirmed at 7 dpi (10 day post dsRNA injection) (Fig 5A). Finally, viral burden was also measured at 7 day post infection (10 day post dsRNA injection), observing a significant reduction in DENV titers (5B). These results show that AeSNAP and ATPase proteins are involved in DENV dissemination control in the *Aedes aegypti* mosquito vector.

**Figure 4.**
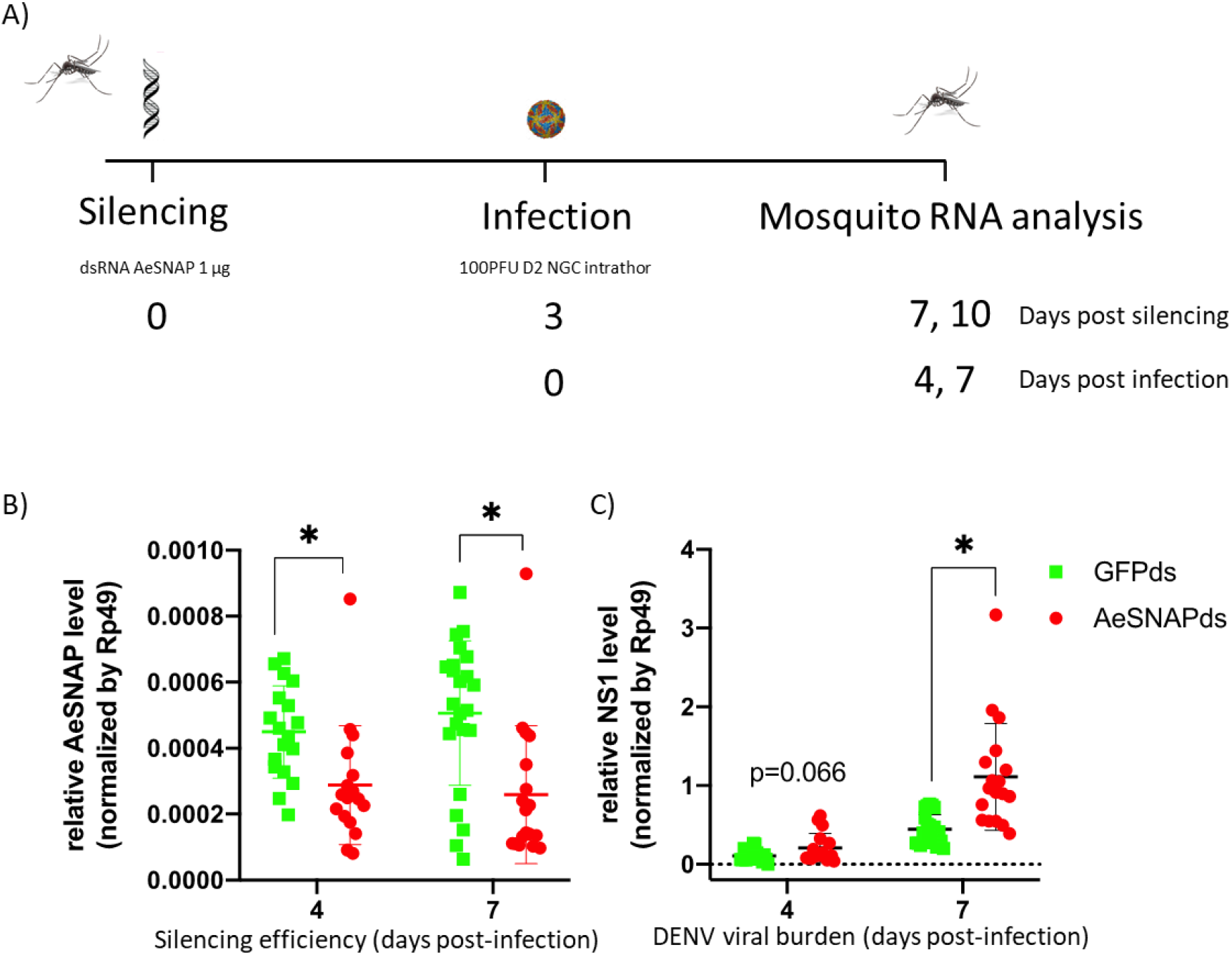
Dissemination analysis of DENV2 in AeSNAP dsRNA-knockdown mosquitoes. (A) Scheme of the strategy for dissemination studies in the *Aedes* mosquito. *A. aegypti* mosquitoes were intrathoracically injected with AeSNAP dsRNA, and at 72h, they were infected with 100PFU of DENV2 using the same route. Silencing efficacy and viral burden were evaluated at 4- and 7-day post infection. B) AeSNAP silencing efficacy. (C) DENV2 viral load recovered from DENV2 infected *A. aegypti* mosquitoes. AeSNAP and DENV2 RNA levels were analyzed by qRT-PCR and normalized to the levels of *Rp49*. Asterisks represent significant difference between samples, calculated by Mann-Whitney non-parametric test (p≤0.05).

**Figure 5.**
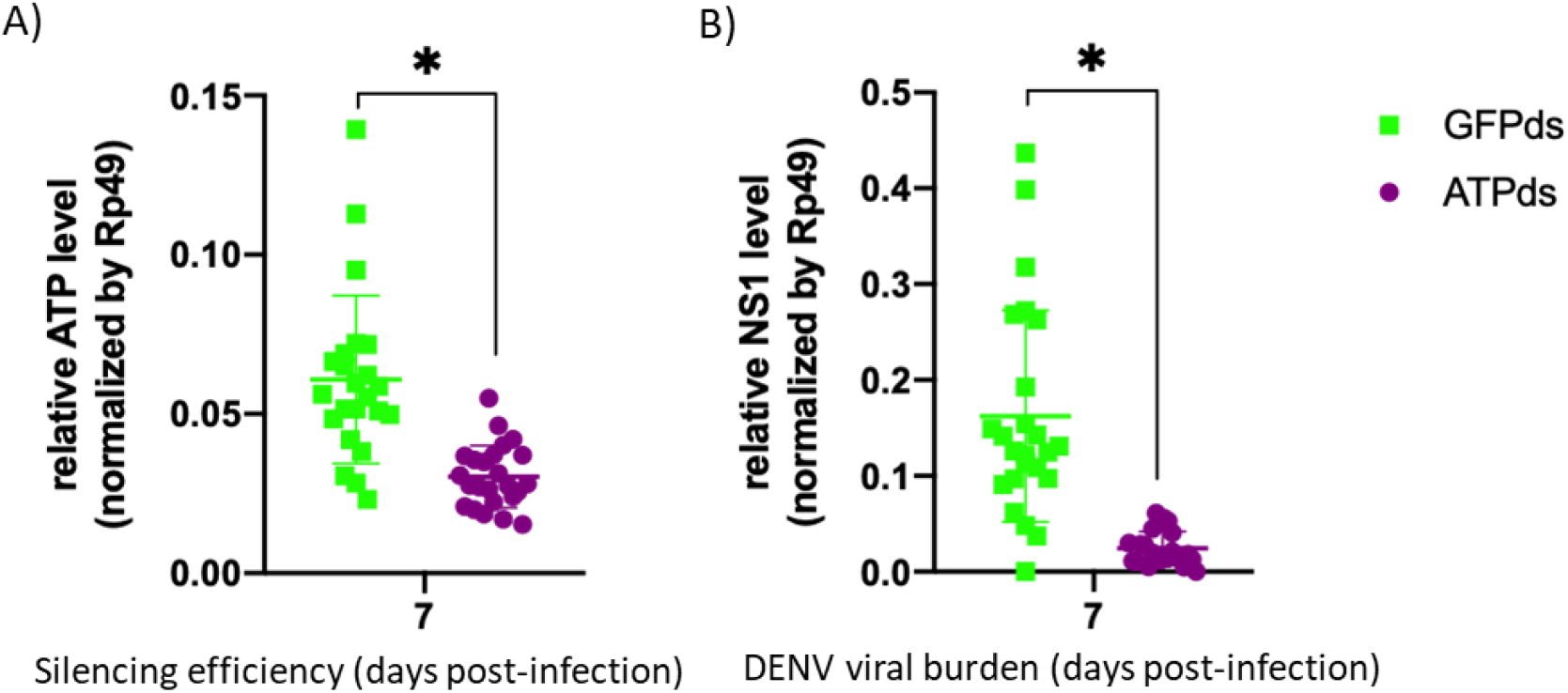
Dissemination analysis of DENV2 in ATPase dsRNA-knockdown mosquitoes. A) ATPase silencing efficacy. B) DENV2 viral load recovered from DENV2 infected *A. aegypti* mosquitoes. ATPase and DENV2 RNA levels were analyzed by qRT-PCR and normalized to the levels of *Rp49*. Asterisks represent significant difference between samples, calculated by Mann-Whitney non-parametric test (p≤0.05).

## Discussion

Infectious diseases transmitted by arthropod vectors, especially by mosquitoes, have acquired increasing medical importance over the few last decades. Among arthropod-borne viral infections, Dengue virus (DENV) is the most prevalent: more than 3.9 billion people in over 129 countries are at risk of contracting dengue, with an estimated 96 million symptomatic cases and an estimated 40,000 deaths every year (World Health Organization, Vector borne diseases). Although a DENV vaccine was just recently licensed by the U.S. Food and Drug Administration for the first time ever, it is far from ideal and its use is restricted only to seropositive individuals, given the excess risk of severe dengue in seronegative vaccinees (21), where sub-optimal immunogenicity in first immune response to dengue predisposes them to a higher risk of severe disease when they experience their first natural dengue infection (ADE phenomena).

Other approaches, including insect vector control and blocking pathogen transmission within these vectors, are promising tools to control the spread of DENV (22). To achieve this goal, it is necessary to understand the molecular mechanisms underlying the interactions between DENV and proteins in the *A. aegypti* mosquito. Success in prevention of pathogen transmission will primarily be based on targeting mosquito proteins which confer resistance or facilitates the infection within the vector. To systematically analyze potential mosquito proteins which interact with DENV particles and could have a role during viral infection, we performed a mass spectrometry assay using purified DENV2 particles and *A. aegypti* salivary gland extracts. We identified a set of *A. aegypti* salivary gland proteins which potentially interact with the DENV virions. After this initial screening, we performed studies of silencing expression by RNAi in selected targets found in the mass spectrometry assay, both *in vitro* and *in vivo*. We demonstrated that two of these proteins, a synaptosomal-associated protein (AeSNAP) and a calcium transporter ATPase protein (ATPase) are involved in DENV viral burden regulation *in vivo*.

AeSNAP belongs to the SNAP family, which are implicated in intra-cellular trafficking and controlling a series of vesicle fusion events (23). These proteins are regulators of vesicle trafficking in synaptic transmission (24), and have additional functions in autophagy and other endocytic and exocytic trafficking processes (25, 26). Moreover, the capsid phosphoprotein P of human para influenza virus type 3 (HPIV3) binds SNARE domains in SNAP29 protein, preventing binding of SNAP29 with SYX17, and hindering the formation of the ternary SNARE complex with VAMP8, required for autophagosome degradation (27, 28). SNAP29 binds the non-structural protein 2BC of the enterovirus-A71 (EV-A71), stimulating autophagy for its replication (29). We show here that RNA interference-mediated knock-down of AeSNAP, an *A aegypti* mosquito protein, leads to an increase in the DENV viral burden in an *A. aegypti* cell line at 24 hpi and also in the whole organism *in vivo*. Surprisingly, in the *in vitro* analysis, we observed a significant decrease in the viral burden at early times post infection (6, 9 and 12 hpi), although this early reduction in the viral burden decreased gradually from 6 to 12 hours post-infection. This phenomenon could be explained by AeSNAP acting at multiple stages of the viral life cycle, showing both an antiviral (late) and a proviral (early times) behaviors, as it has been described for viperin or adenosine deaminases acting on RNA proteins (ADARs) (reviewed in (30, 31)), although this fact should be further explored. This increase in the DENV viral burden under AeSNAP knockdown expression is consistent with the study previously mentioned, in which the HPIV3 capsid phosphoprotein P binds SNAP29, blocking autophagosome degradation and increasing virus production (27) and also with other studies focused on SNAREs and viral burden. For example, Ren et. al showed that inhibition of syntaxin 17 expression by specific small interfering RNAs resulted in an elevated amount of intracellular retained viral particles which facilitated the release of HCV virions by impairing of autophagosome-lysosome fusion (32).

We also identified an *A. aegypti* calcium transporter ATPase protein, ATPase, in our mass spectrometry assay. Calcium transporter ATPase proteins of the sarco (endo) plasmic reticulum (SERCA), the plasma membrane (PMCA), and the secretory pathway (SPCA) are crucial for muscle function, calcium cell signaling, calcium transport into secretory vesicles, mitochondrial function, and cell death (33–35). Several viruses regulate host cell calcium concentrations in the cytoplasm and mitochondria, allowing viral gene expression and replication. For instance, a recent study performed in the human HAP-1 cell line revealed how that Measles virus (MV), West Nile virus (WNV), Zika virus (ZIKV), and Chikungunya virus (CHIKV), and also DENV use the host calcium pump secretory pathway calcium ATPase 1 (SPCA1) for calcium loading into the trans Golgi network, activating glycosyl transferases and proteases and then allowing viral maturation and spreading (36). In our study, we found that the knockdown of this calcium transporter ATPase protein strongly reduced DENV burden in both the *A. aegypti* cell line and the *A. aegypti* mosquito, demonstrating a significant positive association between the level of ATPase protein and DENV viral burden. This finding is in line with another study, in which Vero cells treated with the SERCA-specific inhibitor Thapsigargin showed a significantly reduced level of viral replication for Peste des petits ruminants virus (PPRV) and Newcastle disease virus (NDV) (37). Viruses are small intracellular parasites and rely on protein interactions to produce progeny inside host cells and to spread from cell to cell (38).

Understanding virus-host protein interactions in the mosquito vector can shed light on viral replication and resistance mechanisms. Furthermore, it could lead to important clinical translations, including the development of new therapeutic and vaccination strategies. In this study, we used a mass spectrometry screening assay to characterize a diverse group of mosquito proteins that are potentially associated with DENV virions, and characterized two of these, a synaptosomal-associated protein (AeSNAP) and a calcium transporter ATPase (ATPase) protein, in greater detail *in vivo*. We show that AeSNAP participates in DENV infection control, as its inhibition by RNAi led to a higher viral burden, whereas ATPase seems to be required for DENV infection in both the Aag2 mosquito cell line and in the *A. aegypti* mosquito vector. Further studies are needed in order to identify the specific pathways in which these two proteins are involved, and how they are mechanistically involved with DENV regulation, as well as to analyze other candidates (such as Vtype, Ric) described in the mass spectrometry list that were observed to alter DENV viral burden *in vitro* to a lesser degree. Finally, this study suggests that these techniques can be used to examine interactions between other microbes and components of arthropod saliva. The identified components have the potential to serve as targets for preventing pathogen dissemination in the vector or the transmission to the vertebrate host.

## Materials and Methods

### Cell culture and virus production

Two *Aedes spp*. cell lines were used in this study, Aag2 and C6/36 cells. The *A. aegypti* cell line, Aag2 (ATCC, VA), was used for *in vitro* silencing studies described below. Aag2 cells were grown at 30 °C with 5% CO2 in DMEM high glucose media supplemented with 10% heat-inactivated fetal bovine serum (Gibco), 1% penicillin-streptomycin. In addition, the *A. albopictus* cell line, C6/36, was used to grow DENV stocks using the same media. The dengue strain DENV-2 New Guinea C was used. C6/36 cells were infected at an MOI of 1.0. The culture supernatant was harvested 6 days after infection and subjected to a plaque assay to determine the viral titer, using BHK21 clone 15 cells grown at 37 °C in MEM supplemented as described. Virus stock was stored at −80 °C before use.

### Mosquitoes

*A. aegypti* (Orlando strain, obtained from the Connecticut Agricultural Experiment Station) mosquitoes were maintained on 10% sucrose feeders inside a 12- by 12- by 12-in. metal mesh cage (BioQuip; catalog no. 1450B) at 28 °C and ~80% humidity with a 14:10 h light:dark photoperiod. Egg masses were generated via blood meal feeding on naïve 129 mice. All mosquitoes were housed in a warm chamber in a space approved for BSL2 and ACL3 research. Mosquitoes were used in these experiments 2-14 days after emergence.

### Preparation of DENV and Salivary Gland Mixture

Cell-free supernatants were taken from a T-150 flask of DENV2-infected C6/36 cells at 10 dpi and overlaid on top of room-temperature 30% sucrose-PBS. Samples were ultra-centrifuged at 100k x g for 2 hours and supernatant was removed before viral pellet was resuspended in ~2 mL serum-free DMEM media. Resuspended virus was overlaid on a room-temperature 30%/60% sucrose-PBS gradient and ultra-centrifuged at 100k x g for 2 hours. Using a flashlight shone from underneath the virus/sucrose-gradient, a viral band was visualized and ~600 μL was carefully pipetted to a new Eppendorf tube. A small aliquot was removed to determine viral titer. Sucrose-purified virus was split into two aliquots of ~300 μL (~3.9 x10^9^ viral particles in each) and extract from 10 *A. aegypti* salivary glands (SGE) in 10 μL was added to one of the aliquots. Both the DENV and the DENV+SGE were incubated for 1 hour at 30 °C, and then diluted to 4 mL before being overlaid on a room-temperature 30%/60% sucrose-PBS gradient and ultra-centrifuged at 100k x g for 2 hours. Both bands of the DENV and the DENV+SGE were collected as described above and heat inactivated for 10 minutes at 65 °C before being frozen at −80 °C. This entire protocol was repeated for three biological replicates.

### Liquid chromatography and tandem mass spectrometry analysis (LC + MS/MS)

DENV only and DENV combined with salivary gland extract samples were submitted to the Interdisciplinary Center for Proteomics at the Yale University, where they were precipitated and resuspended in PBS before liquid chromatography tandem mass spectrometry (LC + MS/MS). Charge state deconvolution and deisotoping were not performed. All MS/MS samples were analyzed using Mascot. Mascot was set up to search the *Aedes aegypti*_201505 database (selected for *Aedes aegypti*, unknown version, 37,800 entries) assuming the digestion enzyme trypsin. Mascot was searched with a fragment ion mass tolerance of 0.050 Da and a parent ion tolerance of 10.0 PPM. Carbamidomethyl of cysteine was specified in Mascot as a fixed modification. Gln->pyro-Glu of the n-terminus, deamidated of asparagine and glutamine and oxidation of methionine were specified in Mascot as variable modifications. Scaffold (version Scaffold_4.4.8, Proteome Software Inc., Portland, OR) was used to validate MS/MS based peptide and protein identifications. Peptide identifications were accepted if they could be established at greater than 50.0% probability by the Peptide Prophet algorithm with Scaffold delta-mass correction. Protein identifications were accepted if they could be established at greater than 50.0% probability and contained at least 1 identified peptide. Protein probabilities were assigned by the Protein Prophet algorithm. This entire protocol was repeated for three biological replicates.

### dsRNA production, silencing and DENV infection *in vitro*

For gene knockdown, dsRNA was produced from approximately 500 bp coding regions of either *A. Aegypti* candidate analyzed in this study or green fluorescent protein (GFP) as a control. Genes that were found in at least two out of the three independent experiments were selected for further studies (table 1). Briefly, MS gene candidates were cloned in the pMT-BiP-His-V5 vector using cDNAs from Aag2 or salivary gland. PCR was used to produce a DNA template with T7 overhangs that was then used to generate the dsRNA molecules (TranscriptAid T7 High Yield Transcription Kit, ThermoScientific), according to manufacturer’s instructions. Oligos used for making dsRNA are shown in table 2. For *in vitro* studies, the dsRNA molecules were transfected into Aag2 cells (INTERFERin®, Polyplus) according to manufacturer’s instructions. Briefly, 500 ng of dsRNA were added to 5 x 10^5^ cells in a 48 well plate. 48 h post-transfection, silencing level was analyzed, cells were infected with DENV2 at MOI 1.0, and DENV2 viral burden was analyzed at different timepoints.

**Table 2.**
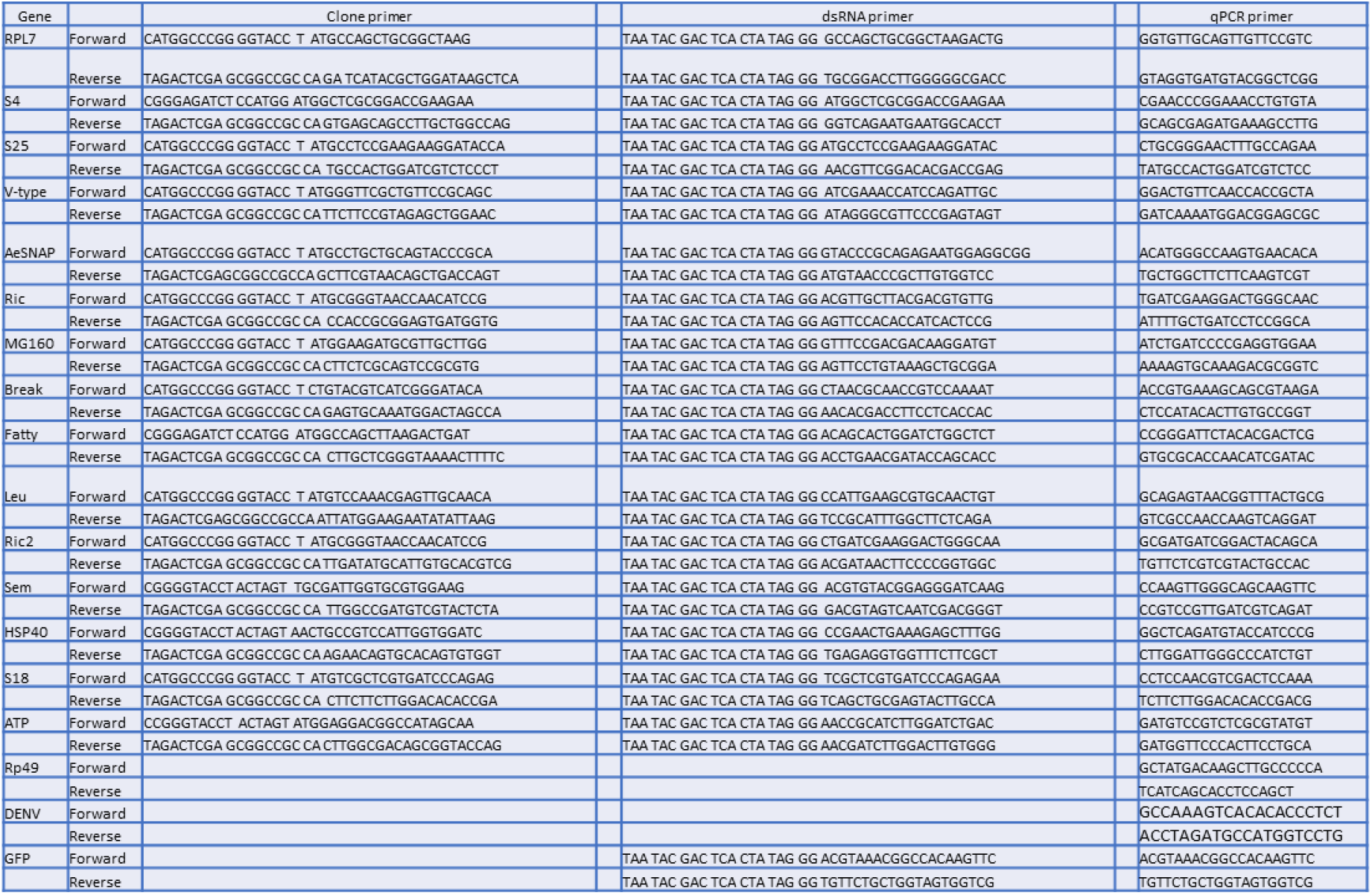
List of primers used for cloning, RNA knockdown and qRT-PCR analysis. Gene candidates found in at least 2 independent experiments were cloned in the plasmid vector. dsRNA molecules were generated for RNAi experiments and the gene expression was analyzed by qRT-PCR.

### *In vivo* mosquito silencing and infection

For dissemination studies, female mosquitoes were injected with dsRNA to silence the individual genes in the salivary glands. As a control, mosquitoes were injected with dsRNA for GFP (dsGFP). Mosquitoes were kept on ice for 15 min, and then transferred to a cold tray to receive an intrathoracic microinjection via the lateral side of the thorax of 1 μg of dsRNA diluted in 138 nl of MQ water, using a Nanoinject II Injector (Drummond Scientific, USA). After injection, the mosquitoes were transferred into cylindrical containers fitted with a nylon mesh on the top and supplied with 10% sucrose solution. 72 hours post dsRNA injection, mosquitoes were infected with DENV2 through intrathoracic injection. Female mosquitoes were immobilized in a cold tray and intrathoracically inoculated with 100 PFU of DENV2 in 138 nl, as previously described. The infected mosquitoes were then dissected on days 4 and 7 after infection to analyze the levels of AeSNAP or ATPase and DENV2 by quantitative reverse transcription PCR (qRT-PCR).

### RNA extraction, cDNA synthesis and qRT-PCR-based assays

Whole mosquito body RNA extractions were performed using TRIzol according to manufacturer’s protocol (Invitrogen, Carlsbad, CA). The RNA was subsequently used for production of a cDNA pool with iSCRIPT (BIORAD). The qRT-PCR assay was done using the iTaq kit according to the manufacturer’s instructions (BioRad). Oligos for the qRT-PCR reactions are shown in table 2. Viral RNA or *Aedes* gene expression was normalized to Rp49 expression. Each sample was tested in quintuplicate for the *in vitro* studies.

### Statistical analysis

GraphPad Prism software was used to perform statistical analysis on all data. Transcription levels of DENV and *Aedes* candidates in mosquito cells, whole mosquito, were normalized using Rp49 housekeeping. Transcription levels were calculated using non-parametric Mann-Whitney Test, as indicated in the figure legends. Asterisk represents P value < 0.05.

### Ethics statement

All experiments were performed in accordance with guidelines from the Guide for the Care and Use of Laboratory Animals (National Institutes of Health). The animal experimental protocols were approved by the Institutional Animal Care and Use Committee (IACUC) at the Yale University School of Medicine (assurance number A3230-01). All infection experiments were performed in an arthropod containment level 3 lab (ACL3) animal facility according to the regulations of Yale University. Every effort was made to minimize animal pain and distress.

## Acknowledgments

This work was supported by NIH grant (AI089992) and (AI127865). E.F. is an investigator supported by the Howard Hughes Medical Institute. The authors also thank Kathleen DePonte for her technical assistance.

